# Antigen properties shape organization of FcεRI aggregates to tune mast cell signaling

**DOI:** 10.1101/2023.08.04.552060

**Authors:** Derek A. Rinaldi, William K. Kanagy, Rachel M. Grattan, Jon Christian David, Hannah C. Kaye, Eric A. Burns, Marelessis Palomino, Shayna R. Lucero, Michael J. Wester, Lydia Tapia, Bruna Jacobson, Keith A. Lidke, Bridget S. Wilson, Diane S. Lidke

## Abstract

Fc receptors containing immunoreceptor tyrosine-based activation motifs (ITAMs) are critical components of the innate immune system that bridge adaptive antibody recognition to cellular effector responses. In allergic responses, the high-affinity IgE receptor, FcεRI, is activated when multivalent antigens crosslink receptor-bound IgE, yet the molecular mechanisms linking antigen structure to signaling output remain incompletely understood. Here, we compare two antigens presenting identical IgE-binding haptens but differing in geometry: the high-valency, heterogeneous DNP-BSA and the defined trivalent antigen DF3. We find that these ligands elicit distinct patterns of degranulation and FcεRI γ-chain phosphorylation, correlating with differences in the recruitment of the inhibitory lipid phosphatase SHIP1. Monte Carlo simulations predicted that each antigen generates receptor aggregates with distinct size, complexity, and inter-receptor spacing. Using direct stochastic optical reconstruction microscopy (dSTORM) and Bayesian Grouping of Localizations (BaGoL) analysis, we directly visualized the nanoscale aggregate geometry and found that DF3 induced smaller, more linear aggregates with tighter receptor spacing than DNP–BSA. Together, our results show that antigen properties, including size, valency, and epitope spacing, modulate FcεRI aggregate architecture and tune the balance of positive and negative signaling to ultimately shape mast cell outcomes.

**Statement of Significance:** Allergic immune responses are initiated when multivalent antigens aggregate IgE-bound FcεRI on mast cells, yet the relationship between antigen structure and signaling strength remains unclear. This study combines biochemical assays, Monte Carlo modeling, and super-resolution imaging to show that allergen properties, specifically valency and nanoscale epitope spacing, govern the geometry of FcεRI aggregates and the balance of activating and inhibitory signals. These findings establish a direct mechanistic link between antigen structure and immune receptor output, providing new insight into how physical antigen features encode mast cell responses.

## Introduction

Crosslinking of IgE-bound FcεRI by multivalent antigen initiates mast cell degranulation and the release of inflammatory mediators that underlie multiple immune responses, including allergy, asthma and anti-parasite immunity (1). FcεRI is a heterotetrametric complex consisting of an α-subunit, a β-subunit and two disulfide-linked γ-subunits. The α-subunit binds circulating IgE with nanomolar affinity (2). Crosslinking of IgE-bound receptor by multivalent antigen leads to recruitment of Src Family Kinases (SFK) that phosphorylate the immunoreceptor tyrosine-based activation motifs (ITAMs) on the β and γ_2_ subunits. Signaling is then propagated by recruitment of SH2-containing downstream signaling molecules to phosphorylated ITAM tyrosines (3–6). The β-subunit is typically considered a negative regulator through SH2-mediated recruitment of phosphatases, such as SHIP1 (7–10), while the γ_2_ subunits primarily drive positive signaling through recruitment of Syk (5,6) which further transduces signaling via activation of the membrane scaffolding protein LAT (11,12). Although positive roles for the β-ITAM in Lyn and Syk binding have also been described (10). The antigens that IgE-FcεRI engages span more than six orders of magnitude in size, from surface arrays on helminths (13) to multivalent pollen proteins (14), food-derived peptides (15,16), and venom components (17). Despite decades of work, a central mechanistic question remains: how do the physical attributes of an antigen translate into a specific FcεRI signaling outcome?

Previous studies have addressed this question using synthetic antigens to show that FcεRI signaling is sensitive to antigen valency, epitope spacing, size, affinity and dose (7,18–28). Baird and colleagues designed rigid DNA rulers to show that there is a strict dependence on antigen valency and size (20–22). Bivalent DNP-DNA triggered degranulation when the spacing was < 5.5 nm but signaling was lost when the distance was increased to just 6.8 nm (22). Increasing the valency to three with a Y-shaped DNA structure supported degranulation out to 10.4 nm spacing (21), highlighting the impact of valency. However, β- and γ-ITAM phosphorylation was still found to decrease as distance increased. The authors concluded that Lyn-mediated ITAM phosphorylation is more efficient when receptors are in close proximity. Using the symmetrically arrayed T4 fibritin foldon, Mahajan *et al*. constructed a structurally defined trivalent antigen, termed DF3 (7). DF3 activation in both RBL-2H3 cells and bone-marrow–derived mast cells (BMMCs) displayed high-dose inhibition, where degranulation peaked at an optimal antigen concentration and then fell off as the dose was further increased. They found that high dose inhibition was a result of increased recruitment of phosphorylated SHIP1 to FcεRI aggregates compared to the optimal degranulation dose. Complementary work with engineered natural allergens is consistent with these findings. For example, Wilson and colleagues showed that progressive digestion of shrimp tropomyosin revealed a lower size limit (~60 aa) for productive signaling (18) that is a result of reducing high valency intact protein to monomeric peptides. Gieras et al generated myoglobin constructs bearing one to four Phl p 1 epitopes and showed valency-dependent severity of anaphylaxis (2,19). Notably, using negative stain electron microscopy, Gieras et al showed that the IgE-Phl p 1 immune complexes increased in complexity with epitope valency. While antigen multivalency is a well-accepted requirement for FcεRI activation, we and others have shown that monovalent ligand presented in a fluid supported lipid bilayer (SLB) can support mast cell secretion and cytokine production (24–26). Suzuki et al compared FcεRI signaling induced by low (NP)- and high (DNP)-affinity antigens presented on a SLB and found that NP elicited reduced degranulation and cytokine production but had enhanced chemokine secretion (26). These differences were attributed to the ability of NP to preferentially recruit the SFK Fgr over Lyn and the scaffold NTAL over LAT. Collectively, these studies establish that antigen properties act to tune FcεRI signaling by modulating the presence of positive and negative signaling molecules in the aggregate.

To better understand the underlying mechanisms connecting antigen-induced aggregate organization to differential signaling, we compared the response of the model mast cell line Rat Basophilic Leukemia-2H3 (RBL) cells primed with DNP-specific IgE (29) to either multivalent DNP-BSA (valency ~ 14) or trivalent DF3. As might be expected, the higher-valency DNP-BSA was markedly more effective than DF3 at inducing FcεRI phosphorylation and mast cell degranulation. Unlike DNP-BSA, which showed a sustained degranulation across a broad range of doses, DF3 exhibited a bell-shaped dose-response curve characteristic of high-dose inhibition. Notably, DNP-BSA triggers robust signaling even in the absence of Lyn kinase, whereas DF3 fails to support degranulation without Lyn. The ability of FcεRI aggregates to recruit SHIP1 was found to be greater with DF3 activation, contributing to the reduced signaling. Structural modeling of IgE-FcεRIα aggregation predicted that DF3 would form smaller, less complex aggregates than DNP-BSA. These predictions were experimentally validated using dSTORM super-resolution imaging and a novel image analysis algorithm, Bayesian Grouping of Localizations (BaGoL) (30) to map the nanoscale organization of FcεRI aggregates. We found differences in the relative arrangement of DNP-BSA and DF3 aggregates, with DNP-BSA aggregates being larger and more complex than DF3 aggregates. Together, these results show that the structural properties of an antigen can directly influence mast cell signaling through spatial organization of FcεRI aggregate architecture.

## Materials and Methods

### Antibodies, Antigens, and Reagents

Modified Eagle’s media (MEM) was purchased from Cytiva (SH30024.01). G418 Disulfate was obtained from Caisson Labs (G030-5GM). Heat inactivated FBS was from R&D Systems (S12450H). L-Glutamine (25030-081), Pen Strep (15140-122), and DNP-BSA (A23018) were purchased from Thermo Fisher Scientific. For unit conversion, 14 nM DNP-BSA is ~ 1 μg/mL. Custom DF3 peptide was synthesized by and purchased from Anaspec, as described in Mahajan et al (7). Anti-H1-DNP-ε-206 IgE was affinity-purified from ascites (Covance, Denver, PA) according to the methods of Liu et al (29). Alexa Fluor 647-(Thermo Fisher Scientific; A-20006), and CF640R-(Biotium; 92108) labeled IgE were prepared using NHS Ester conjugation with a dye/protein ratio of 0.9 and 1.1, respectively. The antibodies used in this study include: anti-pY PY20/99 cocktail (Santa Cruz; sc-508, sc-7020), goat anti-mouse-HRP (Santa Cruz; sc-2005), goat anti-rat IgE (Abnova; PAB29749), and biotinylated anti-HA Fab fragments (Roche; 12158167001). Fura-2 AM (F1221) and QDot 655 quantum dots (QD; Q10111MP, Q10121MP) were purchased from Thermo Fisher Scientific. HA-QD655 was conjugated as previously described (31). Live-cell RBL experiments were performed in modified Hank’s balanced salt solution (HBSS; 136.89 mM NaCl, 5.37 mM KCl, 0.74 mM KH2PO4, 0.70 mM Na2HPO4, 10 mM Hepes, 0.05% w/v BSA, 5.45 mM glucose, 0.88 mM MgSO4, 1.79 mM CaCl2, and 16.67 mM NaHCO3).

### Quantification of DNP Epitope Number on Antigens

A Nanodrop 2000C spectrophotometer was used to measure the absorbance of DNP-BSA at A280 and A360 (the A_max_ of DNP). The molar concentration of BSA was calculated where M_BSA_ = (A280-(A360*CF_DNP_))/ε_BSA_, where ε_BSA_ = 43,824 cm^−1^M^−1^ is the extinction coefficient for BSA and CF_DNP_ = 0.2915 is the correction factor for the contribution of DNP to the A280 reading. The molar concentration of DNP was calculated by M_DNP_ = A360/ε_DNP_, where ε_DNP_ = 17,500 cm^−1^M^−1^. The degree of labelling was determined by the molar ratio = M_DNP_/M_BSA_ and was calculated to be 14 DNP per BSA.

### Cell Culture

RBL-2H3 (RBL-WT), RBL-HA-HL4.1-FAP-FceR1γ, and CRISPR-edited RBL-Lyn^KO^ and RBL-SHIP1^KO^ cell lines (10,32–35) were cultured in MEM supplemented with 10% heat-inactivated fetal bovine serum, 1% penicillin/streptomycin, and 1% L-glutamine. RBL-HA-HL4.1-FAP-FceR1γ cells were cultured with G418-disulfide. RBL-SHIP1^KO^ cells were transfected with a pcDNA 3.1 SHIP1-mNG plasmid (10) by electroporation with an Amaxa Nucleofector II via program T-020 using Solution L (Lonza; VCA-1005) or Mirus Bio Ingenio (MIR-50114). For live cell microscopy experiments, cells were primed with 1 μg/mL unconjugated or CF640R-IgE and incubated overnight in eight-well chambers (Cellvis; c8-1-n) at a density of 4×10^5^ cells/well. High resolution dSTORM was performed on cells primed with 1 μg/mL Alexa Fluor 647-IgE and seeded at a density of 2×10^5^/well in a 6 well tissue-culture plate (Greiner BioOne; 657-160) on 25 mm round cover glass, #1.5H (Warner Instruments; 64-0715). Cells (5×10^6^) were plated on 100-mm tissue culture dishes and primed overnight with 0.5 μg/mL unconjugated IgE for western blot experiments.

### Monte Carlo Simulation of Antibody Aggregation

Model preparation and creation: All-atom crystal structures are used to create space-filling models of IgE and antigens (7,36). IgE is made from multiple PDB structure files 1OAU (37), 2VWE (38), 1O0V (39), 1F6A (2) and DF3 consists of 1RFO (7,40). The BSA structure is one chain of 3V03 (41). DNP-BSA binding sites: As DNP is grafted on BSA by bioconjugation to lysines, we select 14 lysines on the surface of BSA as haptens. Of these 14, 2 are selected as they are the two furthest apart, 1 is picked as approximately equally distant to the previous 2, and the remaining 11 lysines are picked randomly. All models are loaded into ChimeraX (42,43), converted to an isosurface and exported as OBJ.

Monte Carlo Simulation: *Initialization:* The models of IgE and antigens created in the previous step are incorporated into a Monte Carlo simulation that simulates 2D diffusion on the membrane. IgE molecules are oriented with Fab arms pointing away from the membrane plane, and antigens are rotated so that a plane of 3 haptens is parallel to the membrane patch. For simulations of DF3, this means that all haptens are accessible to bind to IgE. For simulations of DNP-BSA, the hapten plane is made by the two lysines that are furthest apart, and the one that is equally distant to them, thus enhancing accessibility of binding sites. We initialize 14 IgE molecules and 14 antigens randomly placed and rotated within a flat membrane patch 200 nm by 200 nm in size. *Update step*: At each step, molecules and aggregates translate on the membrane surface by randomly updating their 2-dimensional center of mass positions, drawing from a normal distribution with variance equal to 2Dt, where D is the diffusion coefficient and t is the time step (t=10 us). The value of the diffusion coefficient for free molecules is D=0.09 um^2/s (23). For aggregates, the effective diffusion coefficient lowers according to D/n (7), where n is the number of IgEs in the aggregate. Rotations of the rigid molecules is also randomly drawn from a normal distribution, with possible values between zero and 360 degrees. Valid moves are those without molecular collisions and within the simulation boundaries. *Binding and unbinding conditions:* Binding between an antigen and IgE is possible if at least one of the antigen’s unbound haptens and one of the IgE’s unbound binding sites are within a certain threshold (1.7nm). If this condition is true, the two molecules bind. Unbinding occurs with a probability of 10^−6^ for each pair of bound binding sites at each step. *Termination condition:* Simulations are run until it reaches the maximum number of time steps or the system becomes unable to sample feasible states (1000 attempts in a single time step). Analysis: Positions and rotations of the molecules’ center of mass are recorded at each time step. At the end of the simulation we compute: (a) aggregate size, as the number of IgE in an aggregate; (b) aggregate spread: as the average distance from an aggregate centroid and all IgE centers of mass within an aggregate, in nm; (c) distances between neighboring IgE, i.e., between IgEs centers of mass that are bound to the same antigen, in nm; (d) distance between centers of mass of antigen and bound IgE, in nm.

### Degranulation Assays

Cells were primed with 1 μg/mL IgE and seeded at 2×10^5^ cells/well in a 24 well plate overnight. Cells were washed twice with warm HBSS and stimulated with a dose course of DNP-BSA or DF3 in HBSS for 30 min at 37°C. Spontaneous granular release was measured in cells without antigen exposure, and total granular content was extracted with 1% Triton X-100 (Sigma Aldrich; T8787). The β-hexosaminidase released into the supernatant was detected by reaction with 4-Nitrophenyl N-acetyl-β-D-glucosaminide (Sigma Aldrich; N9376) in citrate buffer for 1 hr at 37°C. The reaction was stopped with glycine, and β-hexosaminidase concentration was measured by absorbance as 405 nm, as previously described (36).

### Fura-2 AM Calcium Assay

Live cell calcium mobilization assays were performed as previously described (10). Briefly, cells were labeled with 2 μM Fura-2 AM in complete MEM media for 30 min at room temperature and washed with warm HBSS. At acquisition, multiple cells were centered in a field of view and ratio images were acquired at 35°C using an Olympus IX71 inverted microscope outfitted with a UPLANSAPO 60× NA1.2 water immersion objective coupled to an objective heater (Bioptechs). Cells were imaged for 5 min, with antigen added after 30 s of data acquisition. Ratiometric changes in cytosolic calcium were determined by alternating between 340 nm and 380 nm excitation at 1 Hz with a xenon arc lamp monochromator (Cairn Research OptoScan) and collecting the interleaved Fura-2 AM fluorescence emissions at 510 nm with an Andor iXon 887 EMCCD camera. A custom MATLAB script as described in (44) was used to analyze the ratiometric traces of individual cells for the lag time and rise height of calcium release.

### Immunoprecipitation and Western Blot Analysis

Cells were washed with warm HBSS and stimulated as described. Cells were then rinsed in ice-cold phosphate buffered saline (PBS) and lysed on ice for 30 min with NP-40 lysis buffer (150 mM NaCl, 50 mM Tris-HCl, 1% NP-40) supplemented with 1% Halt Protease and Phosphatase inhibitors (Thermo Fisher Scientific; 78446). Whole cell lysates were cleared by centrifugation at 13,000 x g for 20 min at 4°C. FcεRI was immunoprecipitated by first incubating lysates with the anti-IgE antibody on a rotator at room temperature (RT) for 1 hour, followed by affinity purification with Protein A/G PLUS-Agarose beads (Santa Cruz; sc-2003) at 4°C overnight with rotation. Immunoprecipitates were then washed via centrifugation (2500 x g for 3 minutes at 4°C) three times with NP-40 lysis buffer supplemented with Halt Protease and Phosphatase inhibitors. Samples were boiled in 2x non-reducing sample buffer (Bio-Rad; 1658063). After centrifugation and removal of beads from boiled samples proteins were separated via SDS-PAGE electrophoresis (4-20% gel; Bio-Rad; 4568093) and transferred to nitrocellulose membranes using the iBlot2 system (Life Technologies). Membranes were blocked for 30 min in 3% BSA-0.1% Tween-20-tris buffered saline and probed overnight with primary antibodies at 4°C. HRP-conjugated secondary antibody was used for detection and incubated with membranes for 1 hour at RT. Membranes were imaged on the Odyssey FC2 Imager (Li-COR Biosciences) after incubation with SuperSignal West Pico PLUS chemiluminescent substrate (Thermo Fisher Scientific; 34580). Resulting densitometry images were analyzed with Image Studio Software.

### TIRF and BAMF Analysis

RBL-SHIP1mNG cells were incubated overnight with 1 μg/mL IgE-CF640R. Cells were washed with warm HBSS and treated with indicated antigen for 5 minutes at 37°C while imaging by total internal reflection fluorescence (TIRF). Data was acquired on an inverted IX83 Olympus microscope equipped with a 60×1.6X/1.5 NA oil-immersion objective (UPlanApo; Olympus) held at 37°C with a Bioptechs objective heater controller. The excitation source were a 640 nm/140 mW and 488 nm/100mW diode laser (Olympus Soft Imaging Solution; #00026121, #00026125). Images were captured using a Hamamatsu ORCA-Fusion sCMOS camera (C14440) and Hamamatsu W-View Gemini image splitter (A12801-01). For simultaneous 2-color, live cell acquisition a filter set of Chroma single-bandpass 705/100 nm (394148) and a Semrock 520/28 nm (FF02-520/28) were used with a Semrock single-edge standard epi-fluorescence dichroic beamsplitter (FF555-Di03). Images were acquired at 900 ms exposure time. As previously described (10), Bayesian Multiple-emitter Fitting (BAMF) analysis (45) was used to model the signal as either foreground (IgE or SHIP1-mNG) or background emitters. The foreground images of the IgE were used to find borders of IgE aggregates. SHIP1-mNG localizations within the IgE aggregate borders were identified, and the recruited SHIP1-mNG signal intensity was reported versus the corresponding IgE aggregate intensity.

### Kinetics of Immobilization Assay

Single particle tracking to quantify FcεR1 immobilization in response to antigen binding was performed similar to that described in (46). RBL-HA-HL4.1-FAP-FcεRIγ cells (32) were plated in 8-well chambers and primed overnight with 0.5 μg/mL IgE. Before imaging, cells were incubated for 10 min with 500 pM HA-QD655 in HBSS at 37°C and washed four times. Samples were imaged for 10 s prior to addition of antigen. Data was acquired on an Olympus IX71 inverted microscope equipped with a 60× 1.2 N.A and Andor iXon 887 emCCD camera. Cells were maintained at 34-36°C using an objective heater (Bioptechs). Excitation was with a mercury-arc l using a 436/10 nm BP filter (Chroma) and emission was collected using 655/40 nm filter (Chroma). Images were acquired at 20 frames per second. Analysis was performed as described (46).

### dSTORM Super Resolution Cell Fixation and Imaging

RBL-WT cells were washed with warm HBSS, and stimulated at the indicated concentration of DNP-BSA or DF3 in HBSS for 5 min at 37°C. Cells were briefly washed with PBS and fixed in ice-cold 100% methanol (−20°C) for at least 30 min up to several days. Cells were rehydrated in PBS for 30 min after removal from methanol and subsequently washed with PBS three times. Sample were imaged the same day as rehydration using a standard dSTORM imaging buffer with an enzymatic oxygen scavenging system and primary thiol: 50 mM Tris-HCl, 10 mM NaCl, 10% w/v glucose, 168.8 U/mL glucose oxidase (Sigma; G2133), 1404 U/mL catalase (Sigma; C9322), and 33.33 mM β-Mercapto-ethylamine hydrochloride (Sigma Aldrich; 30078), pH 8.5. Data was acquired using a custom-built microscope (38) equipped with a 1.35 NA silicone oil immersion objective and a sCMOS camera (C11440-22CU; Hamamatsu Photonics) (Olympus UPLSAPO100XS). A 647 nm laser was used for excitation (500 mW 2RU-VFL-P; MPB Communications Inc.) and emission was collected with a 708/75 nm filter (FF01-708/75-25; Semrock). A total of 100,000 frames were collected per cell with an exposure time of 50 ms per frame. Cells were registered in X, Y, and Z coordinates every 5,000 frames (47).

### dSTORM Image Reconstruction

dSTORM images were analyzed and reconstructed with custom-built MATLAB functions that are available as part of the *smite* package (48) (https://github.com/lidkelab). Briefly, for each image frame, sub-regions were selected around local maxima that were above a threshold (200 photons). Each sub-region was fit using a finite pixel Gaussian point spread function using a maximum likelihood estimator (49). Fitted results were rejected using a log-likelihood ratio test (50). The fit precision was estimated using the Cramér-Rao lower-bound values for each parameter. Blinking events that spanned multiple contiguous frames were found and connected using the cost matrix approach described in (51). The resulting coordinates were corrected for sample drift by minimizing pairwise distances as described in (52).

### BaGoL Emitter Fitting Method

Coordinate data from dSTORM imaging was post-processed using the BaGoL (Bayesian Grouping of Localizations) algorithm which has been previously described (30). Briefly, the BaGoL grouping method is implemented in MATLAB as part of the *smite* single molecule analysis package(48). It functions to group coordinates from multiple blinking events into estimates of the true number and locations of the underlying emitters. This process improves the precision of the estimate of the emitter locations and facilitates downstream analysis. The parameters for the BaGoL analysis used for this data set were: Length of burn-in chain 64,000; number of trials 16,000; ROI size 500 nm; SE adjust 5 nm; initial blinking distribution Xi [20,1].

### DBSCAN Cluster Analysis

Analysis of dSTORM FcεRI aggregation data was performed using the density-based DBSCAN algorithm implemented in MATLAB as part of smite single molecule analysis package (48,53,54). Parameters chosen were a maximal distance between neighboring cluster points of ε = 35 nm and a minimal cluster size of three observations. Cluster boundaries were produced with the MATLAB “boundary” function using a default methodology that produced contours halfway between a convex hull and a maximally compact surface enclosing the points. ROIs of size 3.0 μm × 3.0 μm (9.0 μm2) were selected from the set of images from which statistics for the circularity, nearest neighbor distance (NND), and area were collected per cluster. Circularity (sometimes known as compactness) is defined as (π*A) / p2; where A is the area of the cluster and p is the perimeter.

### Statistical Analysis

Analysis of calcium data, single particle tracking and super-resolution images was performed using in-house MATLAB code as described above. All statistical tests for comparisons between groups was performed using Graphpad Prism version 10.3.0, with the exception of the Bernoulli Trial analysis that was performed in Microsoft Excel.

## Results

### Comparison of DNP-BSA and DF3 structure

We began by comparing the structural properties of trivalent DF3 and multivalent DNP-BSA. DNP-BSA is a commonly used allergen mimic known to induce a high degree of receptor crosslinking and robust mast cell responses (46, 47). DNP-BSA is, however, structurally ill-defined. DNP is bioconjugated to surface exposed lysines for which DNP:BSA ratios of up to 25:1 can be achieved and labeling is assumed to be Poisson distributed. Figure 1A shows the surface lysines available for DNP conjugation and demonstrates that hapten spacing can have a broad distribution, with a minimum of 1.2 nm for adjacent lysine residues and a maximum of 7.2 nm. The DNP-BSA used here was determined to have an average valency of 14 (see Methods) and thus presents multiple binding permutations. In contrast, DF3 is a structurally well-defined trivalent antigen with ~ 3.0 nm spacing between DNP haptens (Fig 1D).

**Figure 1.**
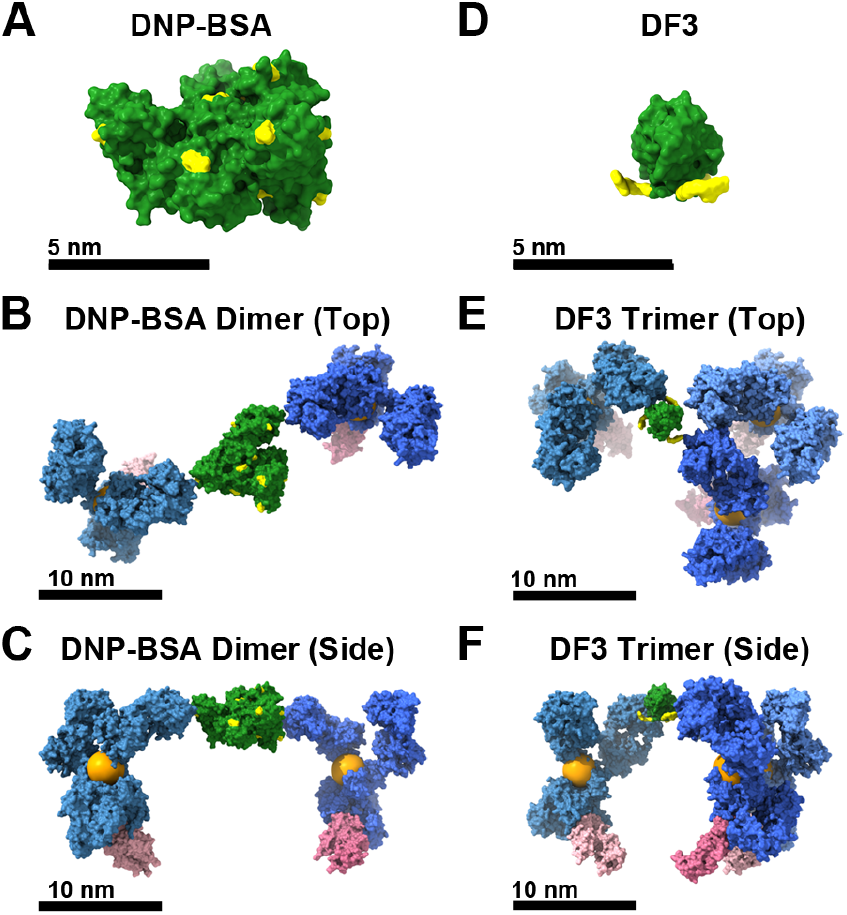
Molecular models of DNP-BSA, DF3 and selected aggregates from Monte Carlo simulation. **(A)** DNP-BSA, with 14 surface Lys selected as possible DNP linking sites shown in yellow. **(B-C)** An IgE dimer crosslinked by DNP-BSA, top and side views. **(D)** DF3 with epitopes shown in yellow. **(E-F)** An IgE trimer crosslinked by DF3, top and side views. Orange spheres in C,F represent the center of mass of IgE from which reported IgE-IgE distances were measured.

To understand how hapten spacing translates into crosslinked IgE/receptor proximity, we used a computational model to estimate the geometry of IgE-FcεRI neighbors when bound to either antigen (Fig 1). This geometric model considers the orientation constraints imposed by binding of IgE to the FcεRI α-subunit at the cell membrane (7). Docking IgE to the farthest separated lysines on DNP-BSA results in a predicted IgE-to-IgE separation of 21.7 nm (Fig 1B-C), as measured from the center of mass of each IgE. However, IgE separation distances will likely include smaller spacings because IgE binding is not limited to the DNP haptens at the extremes. DF3 as a trivalent antigen presents a less variable binding platform. Maximal IgE spacing is smaller with DF3, with separation of 15.2 nm, (Fig 1 E, F). Interestingly, the geometric model predicts that three IgE-FcεRI can simultaneously bind to DF3, but the receptors are organized asymmetrically around the antigen. Figure 1F demonstrates that two of the IgEs are at a maximal spacing of ~15 nm and the third is oriented such that the nearest neighbor is 8.5 nm away. Even with this asymmetry, DF3 binding results in a more controlled distribution of receptor spacing permutations than presented by DNP-BSA. These modeling insights suggest that the two antigens will form aggregates of different structures.

### Antigen properties modulate FcεRI signaling outcomes

We next tested whether the DNP-BSA and DF3 would trigger different signaling by examining the classic outcomes of degranulation and calcium response. Using RBL-2H3 cells primed with anti-DNP IgE (28), we observed the expected robust degranulation with DNP-BSA crosslinking (Figure 1A, gray bars), where the dose response curve is already maximal at 0.01 µg/mL (0.14 nM) and the levels remain high as dose increases. In contrast, DF3 produced degranulation with a bell-shaped dose curve (Fig 1B, gray bars) that peaks at 10 nM DF3. The observed high dose inhibition is consistent with our previous work that first characterized DF3 antigen (6). The differences in cellular response became even more evident when we compared degranulation in cells lacking Lyn, the primary SFK in RBL cells. We recently showed that DNP-BSA can override the need for Lyn, as evidenced by the ability of DNP-BSA to induce degranulation and calcium response in RBL cells where Lyn was knocked out by CRISPR/Cas9 gene editing (RBL-Lyn^KO^;(10)). The ability of FcεRI to signal in the absence of Lyn was attributed to compensation by Lck and/or Src, two SFKs also expressed in RBL cells (see Figure S3 from (10)). Figure 1A confirms that the dose response for DNP-BSA is nearly the same for RBL-WT and RBL-Lyn^KO^ cells. Remarkably, DF3 was not able to support degranulation in RBL-Lyn^KO^ cells at any dose (Fig 1B, red bars).

One of the earliest signaling outcomes after FcεRI crosslinking, and a requirement for degranulation, is an increase in intracellular calcium levels (48). To understand if the loss of DF3-induced degranulation RBL-Lyn^KO^ cells was a result of impaired signal initiation, we next tested the ability of DF3 to induce calcium flux in RBL-WT and RBL-Lyn^KO^ cells. Cells were loaded with the Fura-2 calcium dye and imaged and quantified as described previously (36). We used this method previously to show that DNP-BSA invokes an effective calcium flux in both RBL-WT and RBL-Lyn^KO^ cells, consistent with the observed robust degranulation (10). The heatmaps in Figures 1C & D depict the Fura-2 ratio over time as cells are stimulated with DF3, where each row represents a single cell response. Crosslinking with 10 nM DF3 in RBL-WT cells produced a robust calcium response (Fig 1C). In contrast, the response in RBL-Lyn^KO^ cells with the same antigen and dose was barely detectable (Fig 1D) over unstimulated control cells (Figure S1), consistent with the inability of DF3 to induce degranulation in these cells (Fig 1B, red bars). Notably, only ~ 9% of RBL-Lyn^KO^ cells recorded any detectable calcium response with 10 nM DF3 activation, while DF3-activation of RBL-WT cells resulted in ~ 94% cellular response (Fig 1E). Comparing the magnitude and timing of the cells that did respond shows that RBL-Lyn^KO^ responses are significantly reduced in magnitude (Fig 1F) and slower to initiate (Fig 1G). These results show that different antigens can evoke distinct signaling responses, verifying that mast cell outcomes can be tailored by the specific antigen.

### Antigen properties dictate FcεRIγ phosphorylation levels

We next sought to determine if antigens can differentially modulate the phosphorylation state of the receptor itself. We compared FcεRI γ-subunit phosphorylation across three cell lines RBL-WT, cells lacking the positive regulator Lyn (RBL-Lyn^KO^) described above, and cells lacking the negative regulator SHIP1 (RBL-SHIP1^KO^, described in (10)). Cells were stimulated with either the optimal DF3 dose for degranulation (10 nM) or the closest DNP-BSA dose (14 nM, 1 µg/mL) for 8 min, then lysed and FcεRIγ phosphorylation was probed by western blot. The receptor complex was immunoprecipitated using an anti-IgE antibody and prepared under non-denaturing conditions to retain an intact FcεRI γ-dimer that separates on the gel by phosphorylation state (25, 49). Figure 3A shows a representative western blot, probed for phosphotyrosine (pY), where the FcεRIγ phosphorylation pattern is seen as four bands (1-4 possible phosphotyrosines per dimer; pY1, pY2, pY3, pY4). Notably, the total γ-pY signal for DF3 stimulation is consistently lower than that of DNP-BSA across all cell lines, and DF3 does not induce observable γ-pY banding in the RBL-Lyn^KO^. Therefore, antigen properties directly influence the efficiency of receptor phosphorylation. Figure 3B provides further quantification of the γ-pY banding, revealing that DF3 also induces a less complete phosphorylation profile, not reaching the pY3 and pY4 levels to the same extent as DNP-BSA, in WT cells. Interestingly, the DF3-induced γ-pY distribution is shifted from predominately pY2 in WT cells to the more complete pY3/4 in SHIP1 knockout cells. Together with the results in Figure 2, we found that DF3 signaling is highly dependent on the presence of Lyn and SHIP1 in a way that was not observed for DNP-BSA.

**Figure 2.**
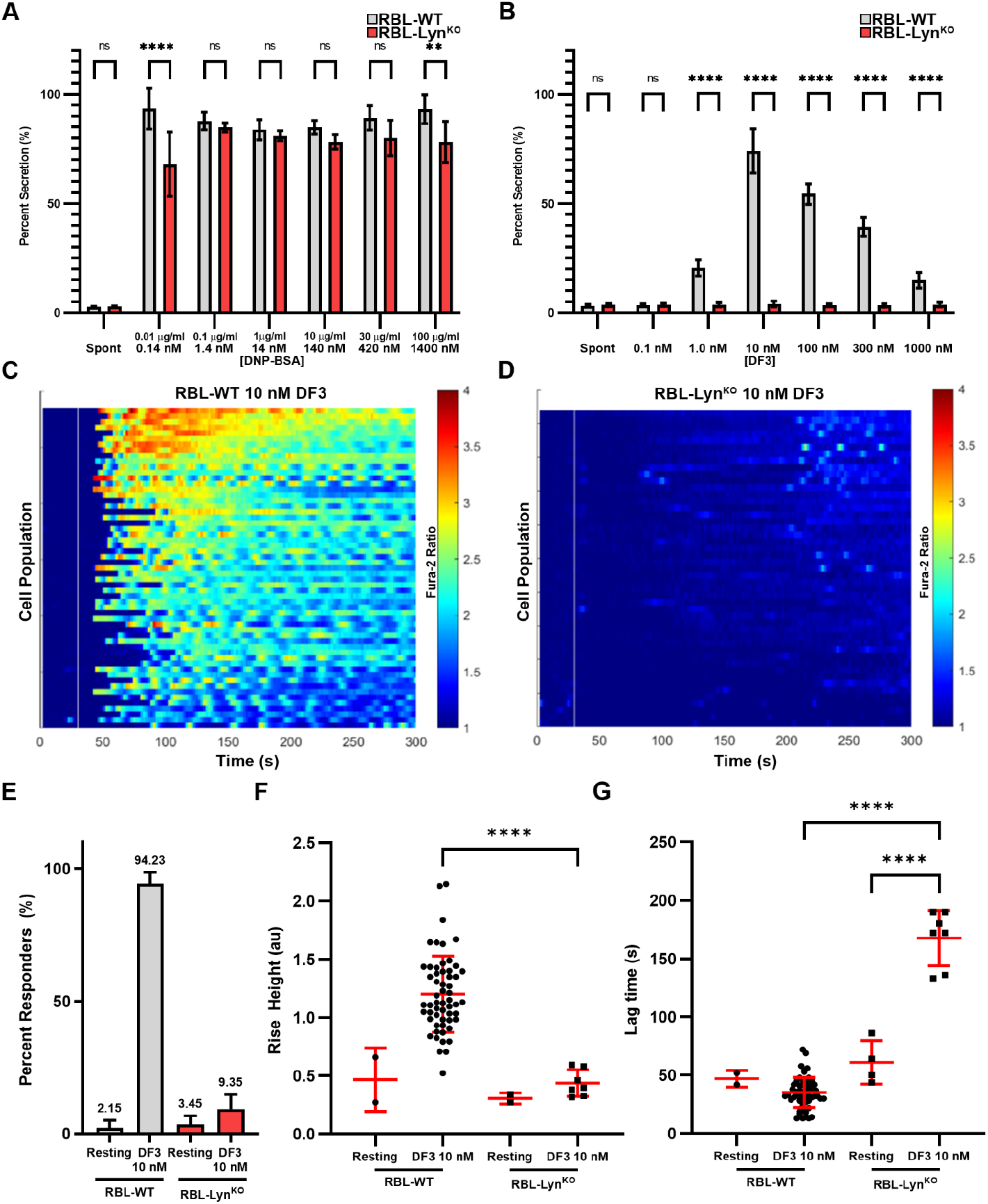
DNP-BSA and DF3 induce differential mast cell outcomes. **A**,**B)** Degranulation results from RBL-WT or -Lyn^KO^ cells. Cells were incubated overnight with 1.0 µg/mL IgE before 30 min of activation with DNP-BSA (A) or DF3 (B) at the listed concentrations. ‘Spot’ = spontaneous release with our antigen. Error bars represent SD of triplicates. n=9 wells across 3 replicate experiments. **C**,**D)** Calcium release in RBL-WT (C) or -Lyn^KO^ (D) cells after antigen activation with 10 nM DF3. Kymographs display the Fura-2 ratio over time on a heatmap, where each row represents single cell; n=104 and n=107, respectively. **E)** The percent of total cells which released detectable calcium after 10 nM DF3 crosslinking, or prior to antigen addition, for both RBL-WT and -Lyn^KO^ cells. Error bars display 95%confidence interval calculated by Bernoulli trial analysis. **F**,**G)** Quantification of the initial magnitude of calcium released (Rise Height, F) and the delay between antigen addition and calcium flux (Lag Time, G). ** p<0.01, *** p<0.001, **** p<0.0001, Two-way ANOVA with Sidak’s multiple comparison test.

**Figure 3.**
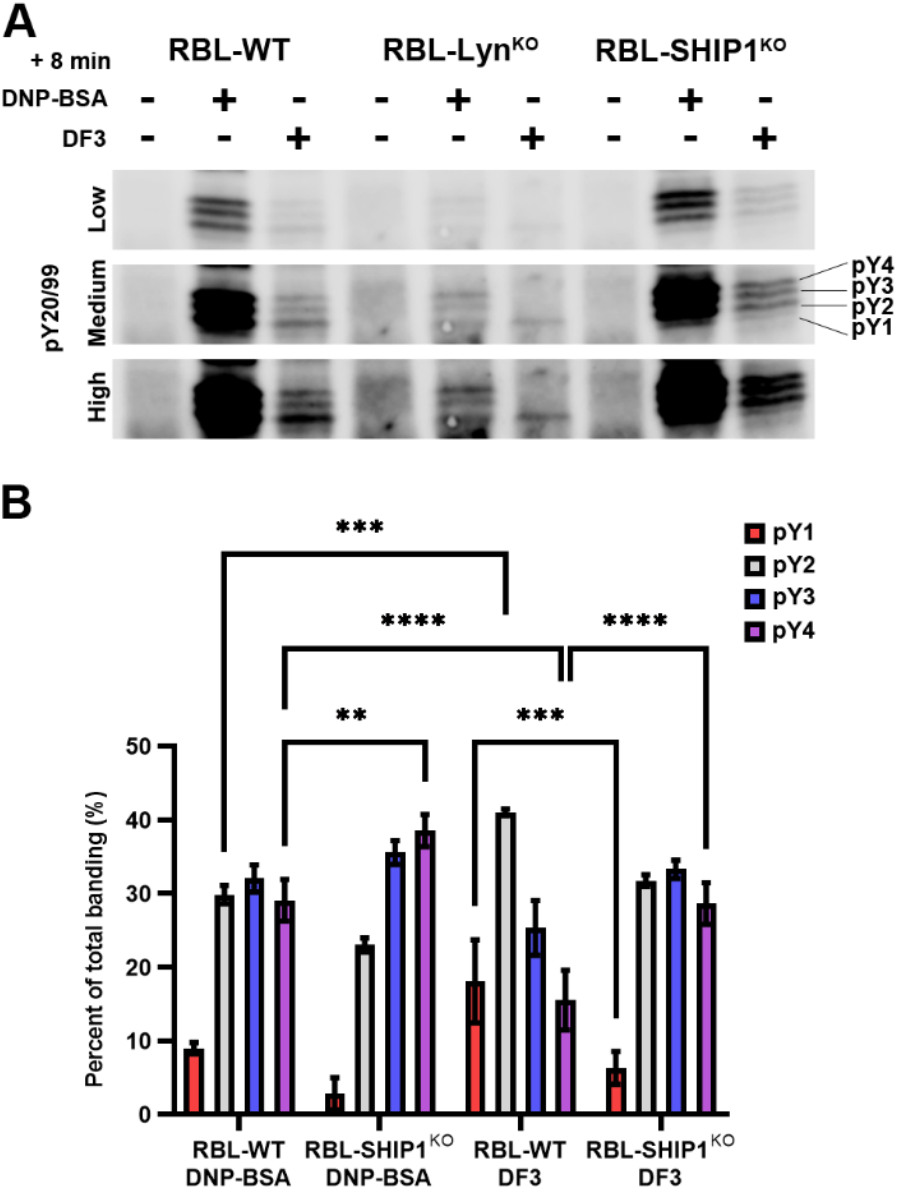
FcεRIγ phosphorylation is stronger and more complete with DNP-BSA than DF3. **A)** Whole cell lysates were collected from RBL-WT, -LynKO or -SHIP1^KO^ cells at rest or after DNP-BSA (14nM) or DF3 (10nM) crosslinking for 8 min. Total tyrosine phosphorylation banding was visualized via western blotting. Low (top), medium (middle) and high (bottom) exposure images are shown to visualize conditions with lower phosphorylation levels. **B)** Densitometry quantification of immunoblots described in (A). Bars represent the percentage contribution of each γ-ITAM-dimer phosphorylation state to the total FcεRIγ activation profile. n=3 independent experiments, **** P < 0.0001, *** P < 0.001, Two-way ANOVA with Tukey’s multiple comparison’s test.

### DF3-induced high dose inhibition is related to enhanced SHIP1 recruitment

Mahajan *et al*. previously showed that the knockdown of SHIP1 using siRNA resulted in an increase in degranulation at all doses of DF3, but the high dose inhibition remained (6). Here, we found that complete knockout of SHIP1 by CRISPR/Cas9 did not alter the DNP-BSA dose response (Fig 4A) yet abrogated the DF3 high-dose inhibition (Fig 4B). This result reveals that the activity of SHIP1 is more sensitive to DF3-induced aggregation than that of DNP-BSA. To understand if distinct antigens can modulate SHIP1 recruitment, we expressed SHIP1-mNG in RBL-SHIP1^KO^ cells and quantified the recruitment of SHIP1-mNG to IgE-CF640R-labeled FcεRI aggregates. Cells were imaged using Total Internal Reflection Fluorescence (TIRF) microscopy after 5 min of crosslinking at the highest antigen dose (Fig 4C, D), in order to assess the point where DNP-BSA and DF3 show the most different outcome. Figure 4E plots the IgE-CF640R intensity in an individual aggregate against the intensity of SHIP1-mNG recruited to that same aggregate, using the Bayesian Multiple-emitter Fitting (BAMF) method as described previously (10,45). At this high dose of antigen, the distribution of FcεRI aggregate intensities was similar between antigens, suggesting that aggregate size alone is not the explanation for the observed difference in SHIP1 recruitment. We found that the amount of SHIP1 recruited to DF3 aggregates of any size (Fig 4E, blue) was consistently greater than that for comparably sized DNP-BSA aggregates (Fig 4E, red).

**Figure 4.**
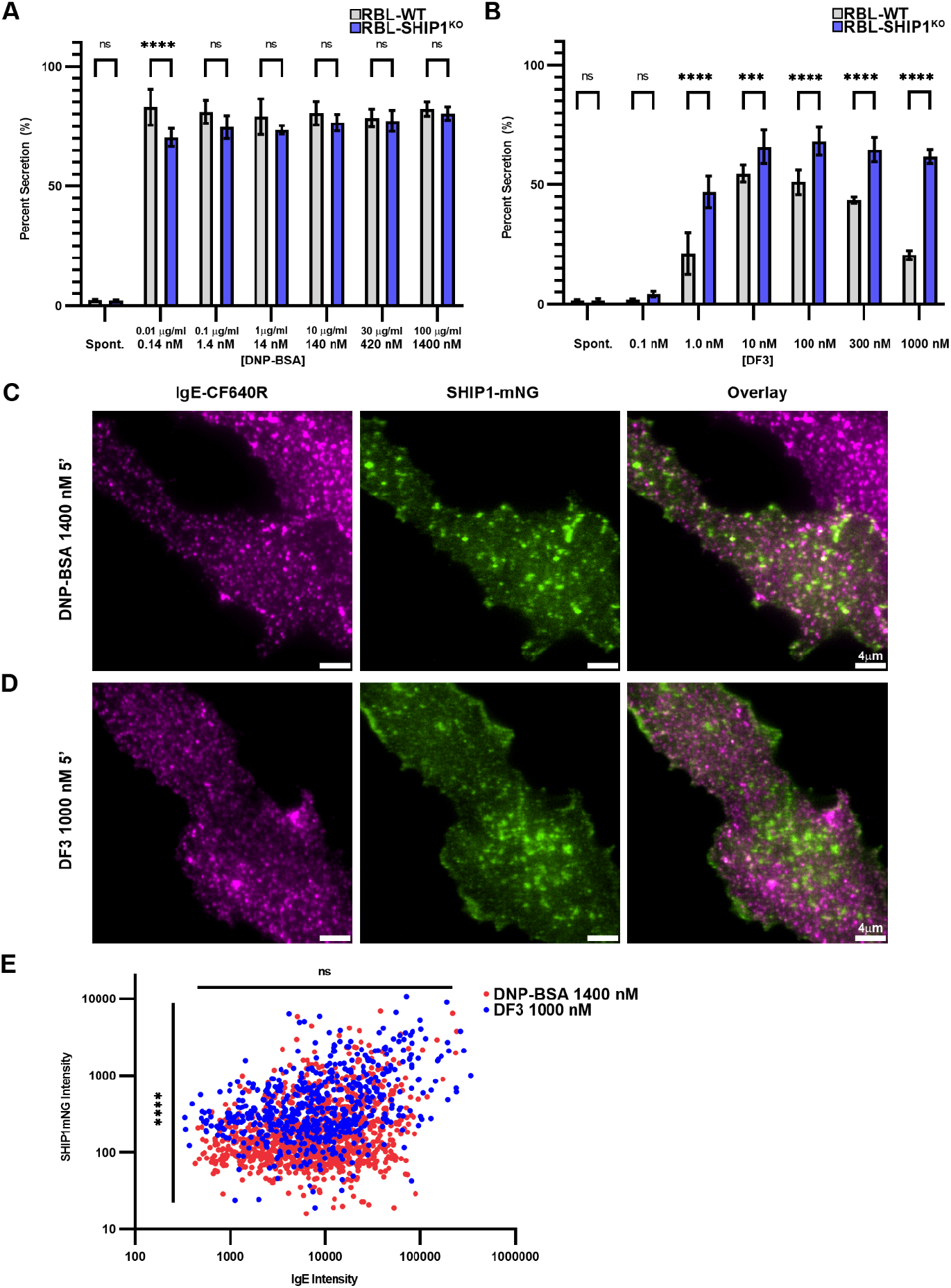
DF3-induced high-dose inhibition is a result of SHIP1 recruitment to FcεRI clusters. **A**,**B)** Degranulation results from RBL-WT or -SHIP1^KO^ cells. Cells were incubated overnight with 1.0 µg/mL IgE before 30 min of activation with DNP-BSA (A) or DF3 (B) at the listed concentrations. Error bars represent SD of triplicates. n=9 wells across 3 replicate experiments. ** p<0.01, *** p<0.001, **** p<0.0001, Two-way ANOVA with Sidak’s multiple comparison test. **C**,**D)** Representative images of RBL-SHIP1^KO^ cells reconstituted with SHIP1-mNG (green) and primed overnight with 1.0 µg/mL IgE-CF640R (magenta), acquired in TIRF. Cells were maintained at 37^°^C during imaging and activated with 1400 nM DNP-BSA (C) or 1000 nM DF3 (D) for 5 minutes. **E)** Quantification of the amount of SHIP1-mNG recruited to individual FcεRI aggregates, plotted as the IgE-CF640R intensity versus SHIP1-mNG intensity per aggregate (see BAMF Image Analysis). n= 6 cells/598 aggregates (DF3) and 1088 aggregates (DNP-BSA). **** p<0.0001, Welch’s unpaired t-test comparing SHIP1-mNG intensities and IgE intensities between antigenic conditions, respectively.

### DNP-BSA and DF3 aggregates have similar aggregation kinetics

We have shown so far that DNP-BSA and DF3 evoke dramatically different signaling outcomes that are associated with differential receptor phosphorylation and SHIP1 recruitment. However, the underlying biophysical mechanisms directing these differences were not yet clear. We next considered whether the binding kinetics of the two antigens may differ. Previous work has suggested that the DNP epitopes on DNP-BSA are not immediately available and aggregation proceeds with a slower kinetics after antigen binding (55). In our own work, we have used single particle tracking (SPT) of quantum dot (QD)-IgE bound receptors to show that receptor diffusion is slowed rapidly upon DNP-BSA binding, and that this immobility is a result of receptor aggregation (23,46). We performed similar SPT experiments, but now using RBL cells expressing an HA-tagged ^γ^-subunit (32). This allows for tracking of individual receptors, without modification of the IgE, by labeling FcεRI γ-subunits with QD-coupled anti-HA Fab fragments (31). To capture the time course of immobilization, antigen was added at 10 s after the initiation of the time series and imaging continued to 50 s. From these movies, we calculated the average diffusion coefficient over a 10-frame sliding window that estimates receptor diffusion before, during and after crosslinking (Fig 5). Consistent with our previous work (23), we found that addition of 14 nM (1 µg/mL) DNP-BSA leads to rapid receptor immobilization, occurring with a rate constant τ = 1.32 s. 10 nM DF3 rapidly induced receptor immobilization on a similar time scale (τ = 1.69 s) as DNP-BSA. Therefore, the observed signaling differences are not likely due to a difference in binding kinetics.

**Figure 5.**
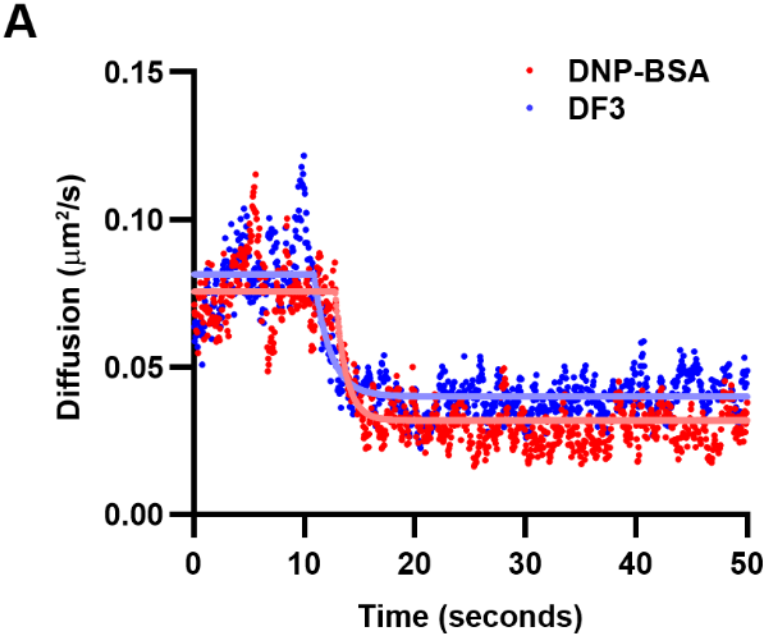
Antigen-induced immobilization kinetics. Plot represents the change in diffusion coefficient over time. Cells were treated with 10 nM DF3 or 14 nM DNP-BSA at 10 s after the start of the time series. Each trace is an average of 8 individual cells. Fits to obtain the rate constant are shown in solid lines.

### Geometric Modeling reveals differences in aggregates formed by DNP-BSA versus DF3

Finding that the antigens form aggregates with similar kinetics, we postulated that it may be the nanoscale organization of the aggregates that direct the differential signaling. To explore this possibility, we first used Monte Carlo simulations to estimate the geometry of IgE-FcεRI aggregates. The model builds on the structures in Figure 1, with the IgE-FcεRIα orientation constrained on the cell surface and incorporates experimentally determined diffusion coefficients (23). The simulation begins with all receptors in the free (no antigen bound) state, and as the simulation progresses the transition to antigen bound (singleton) and the formation of aggregates (short linear chains or more complex, branching aggregates) is monitored (Fig 6A,B). Simulation results indicate that multi-valent DNP-BSA bound to bivalent IgE primarily generates short linear chains, but also induces a considerable fraction of complex aggregates (Fig. 6A). While trivalent DF3 also creates linear chains, complex aggregates are not prevalent (Fig. 6B). Figure 6C shows an example of a branched, complex aggregate formed by DNP-BSA. Figure 6D shows a linear chain of receptors crosslinked by DF3. We can measure aggregate size from the simulations by comparing either the number of receptors per aggregate (Fig. 6C) or the aggregate spread, (Fig. 6D). Our results show that, while both antigens have a peak size of 3 receptors per aggregate, DNP-BSA has the potential to generate larger aggregates (>5 receptors) than DF3 (Fig. 6C). Consistent with this, the aggregate spread, measured from the center of mass of the aggregate, is significantly larger for DNP-BSA than DF3 (Fig. 6D).

**Figure 6.**
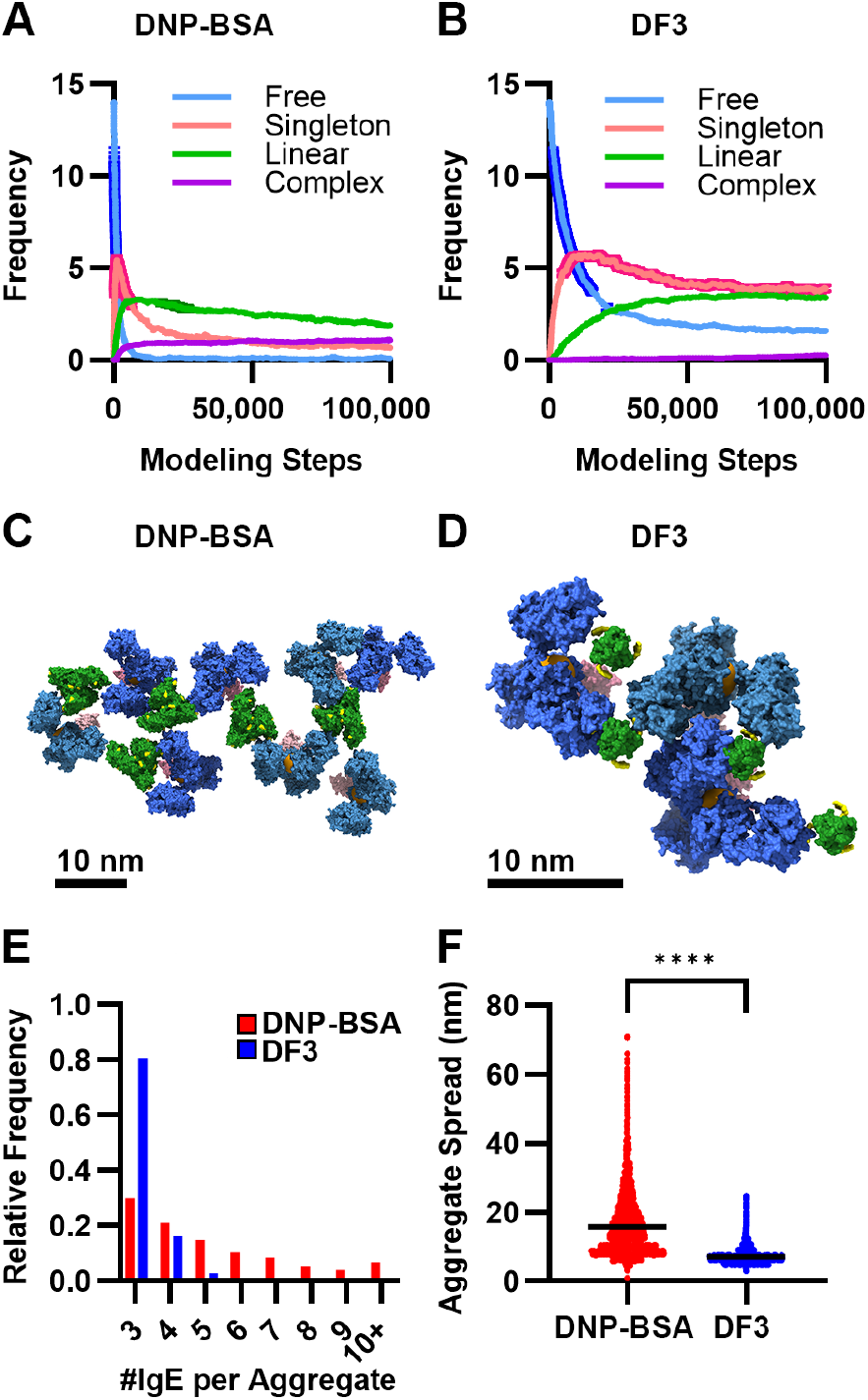
Aggregation dynamics in Monte Carlo simulation. **(A-B)** Types of aggregates that are formed during simulation of IgE-FcεRI^α^ being crosslinked by DNP-BSA **(A)** or DF3 **(B)**. Free (blue line) corresponds to no IgE bound. Singleton (pink line) is IgE bound to allergen, but not crosslinked to another receptor. Linear (green line) is crosslinked IgE forming a linear chain. Complex (purple line) is crosslinked IgE forming a branched aggregate structure. **(C)** Example complex aggregate from DNP-BSA crosslinking simulation (IgE (blue); FcεRI^α^ (pink); antigen (green); binding site (yellow). **(D)** Example linear chain from DF3 crosslinking simulation. **(E)** Aggregate occupancy at the end of the simulation, showing larger aggregates formed by DNP-BSA (red). **(F)** Distribution of aggregate spread, i.e., the average distance from the center of the aggregate to each IgE, indicating that DNP-BSA forms larger and more varied aggregates.

### Signaling efficiency is correlated with aggregate geometry

The Monte Carlo simulations suggested that DNP-BSA and DF3 form aggregates with distinct size and organization. However, it has been challenging to experimentally quantify aggregate geometry on the plasma membrane due to their small size, which is typically below the diffraction limit of light. Super-resolution imaging, such as single molecule localization microscopy (SMLM) techniques, provides sub-diffraction information, but the repeated blinking of fluorophores has made it difficult to identify discrete single emitters (50, 51). Recently, Fazel et al developed an analysis algorithm that increases the precision obtained in SMLM by grouping repeat localizations from the same emitter (29). This is achieved using a Bayesian inference method to perform a weighted average over all possible groupings, ultimately finding the most likely number and positions of the emitters that generated the observed localizations. This algorithm, termed Bayesian Grouping of Localizations (BaGoL), results in increased localization precision (here, the median ranged from 4.7 to 5.7 nm over all conditions) and enables the identification of individual emitters that can be used for further analysis (for complete description of workflow, see Fig. S2). We used dSTORM imaging of IgE-AF647-labeled FcεRI (36, 52) combined with BaGoL analysis to map the lateral organization of individual FcεRI aggregates. Figure 7A-C shows representative BaGoL images and DBSCAN (43) aggregate cluster contours for resting cells, DNP-BSA activated cells or DF3 activated cells. Signaling studies above compared the DF3 maximal dose to the closest DNP-BSA dose from our standard dose curve. For super-resolution imaging of aggregates, we decided to compare 10 nM DF3, where signaling is maximal, with either 10 nM DNP-BSA to match the antigen dose or 2.15 nM DNP-BSA to have equivalent DNP haptens present. Notably, these two doses show similar degranulation to 14 nM DNP-BSA (Fig. S2). From the images, is it already apparent that the lateral receptor distribution is changed upon antigen binding, with DNP-BSA generating larger aggregates than DF3. The Hopkin’s statistic provides a statistical measure of clustering (56) and is calculated for each selected ROI. Plotting the Hopkin’s statistic in Figure 7D for the post-BaGoL images confirms that receptors are nearly randomly distributed in resting cells (value ~ 0.5) and become aggregated (value > 0.5) after antigen binding. While each antigen did induce aggregation, the respective aggregate geometry was found to be different. DF3-induced aggregates were smaller in area compared to DNP-BSA (Fig 7E). Both DNP-BSA and DF3-created aggregates with a broad distribution of shapes, as measured by compactness, ranging from more linear (close to 0) to nearly round (close to 1) (Fig 7F). Yet, the DF3 distribution is shifted towards lower values, indicating a prevalence of linear chains. Comparison of the number of receptors per aggregate showed that DNP-BSA aggregates contain more receptors than for DF3 (Fig 7G). Finally, we calculated the nearest neighbor distances (NND) between receptors within aggregates and found that the receptors are more closely packed in DF3 aggregates (Fig 7H). The most common NND for DF3 is 6.91 nm, while both DNP-BSA doses are shifted to ~8.5 nm. These results support the idea that FcεRI activation is sensitive to nanoscale differences in aggregate geometry that is itself due to specific antigen structural properties.

**Figure 7.**
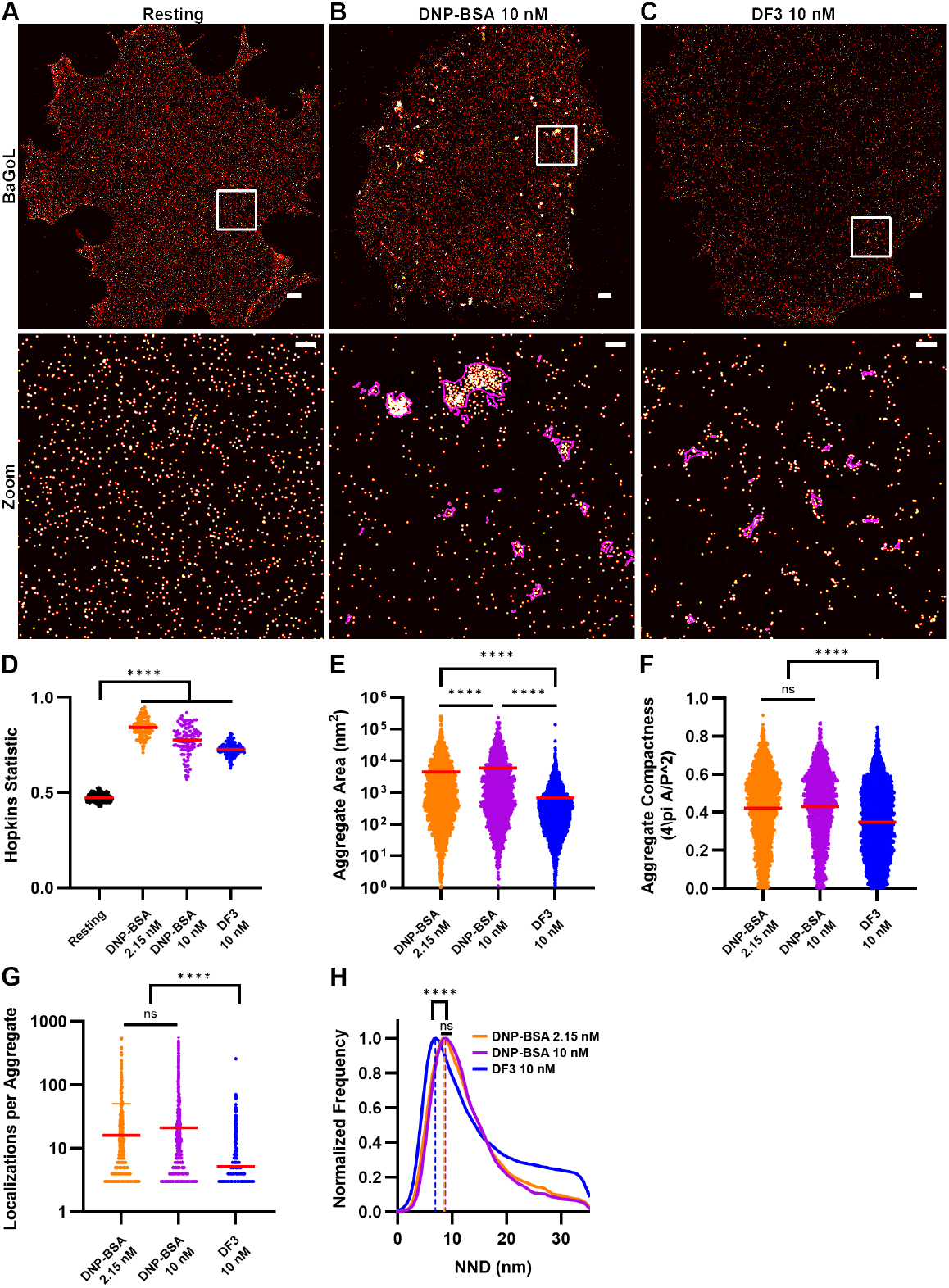
Nanoscale organization of FcεRI aggregates. **A-C)** Representative super-resolution images of resting **(A)**, 10 nM DNP-BSA **(B)** and 10 nM DF3 **(C)** aggregates after 5 min of crosslinking. Top: Single cell images of BaGoL-collapsed dSTORM localizations. Scale bar, 1 µm. Bottom: Magnification of ROI (white box) in image above, where aggregates identified by DBSCAN are outlined in magenta contours. Scale bar, 50 nm. **D-H)** Quantification of aggregate properties from DBSCAN analysis. **D)** Hopkins Statistic indicates that receptors are nearly randomly distributed in resting cells (value ~0.5). Aggregation is seen as the value increase towards 1, with DF3 showing less aggregation than DNP-BSA. Each dot represents the value from a single ROI. **E)** Aggregate area is larger for DNP-BSA crosslinking. **F)** Measurement of aggregate compactness shows that DF3 aggregates are more linear in shape, with values closer to 0. **G)** DNP-BSA aggregates have more receptors than DF3 aggregates. **H)** Histogram of nearest neighbor distance (NND) shows that receptor-to-receptor spacing is closer in DF3 aggregates. Vertical lines indicate the mode for each condition: 6.91 nm for DF3, 8.51 nm for 2.15 nM DNP-BSA, 8.67 nm for 10 nM DNP-BSA. **** P < 0.0001, analyzed by One-way Anova with Kruskal-Wallis Test with Dunn’s multiple comparisons (D-H). N = 3,271 aggregates for 2.15 nM DNP-BSA, 2,205 for 10 nM DNP-BSA and 6,932 for DF3 (E-G) or n = 52,527 calculated receptor spacings for 2.15 nM DNP-BSA, 46,529 for 10 nM DNP-BSA and 35,941 for DF3 (H), acquired over at least 10 cells for each condition. (D-G) The mean of each condition is indicated by a horizontal red bar.

## Discussion

The multivalent structure of allergens is a key regulator of FcεRI clustering and signal initiation. Previous studies have examined how antigen valency and epitope spacing impacts mast cell signaling, with the general trend suggesting that higher valency antigen (3+ epitopes) and close hapten proximity (< 10 nm) produce a stronger activation response (18,21,26). In this study, we sought to better understand how the molecular mechanisms governing FcεRI signaling are tailored in response to specific structural properties of the antigen. We compared two antigens, DNP-BSA and DF3, that present the same DNP hapten but differ in size, valency and epitope spacing. Modeling of each antigen bound to IgE-FcεRIα indicated that the spatial constraints of the antigen are translated into unique receptor aggregate geometry. Degranulation assays showed that each antigen evoked a distinct response, with DNP-BSA causing robust degranulation at all doses and DF3 exhibiting high-dose inhibition. Calcium signaling and FcεRI γ-phosphorylation patterns also differed between antigens.

To understand the underlying mechanisms of the observed differential signaling, we altered the balance of positive and negative signaling by removing either Lyn or SHIP1 via CRISPR/Cas-9 knockout. We found that DF3-induced degranulation is highly sensitive to the presence of Lyn and SHIP1, while DNP-BSA degranulation is largely unaltered in RBL-Lyn^KO^ and -SHIP1^KO^ cells. Despite the robust DNP-BSA-induced degranulation in the knockout cell lines, an influence of SHIP1 and Lyn was seen at the level of FcεRI γ-phosphorylation patterns. We showed previously that DNP-BSA crosslinking overcomes the loss of Lyn by recruiting a compensating SFK that can phosphorylate the γ-ITAM, albeit incompletely (10). We further showed that Syk can bind to γ-ITAMs dimers with mono-phosphorylated C-terminal tyrosines and further enhance ITAM phosphorylation in a feedforward manner, facilitating robust degranulation in the absence of Lyn (10,33). This flexibility in Syk-SH2 binding modes may explain how DF3 can elicit a degranulation response in RBL-WT cells even though the γ-phosphorylation profile is weak and incomplete. On the other hand, we found here that DF3 cannot induce receptor phosphorylation in the RBL-Lyn^KO^ cells at any dose, suggesting that DF3 aggregates cannot recruit other SFKs to the extent needed to overcome negative signaling. Consistent with a role for SHIP1 in high dose inhibition, DF3 aggregates were found to recruit more SHIP1 than DNP-BSA aggregates at high dose. In addition, the DF3 degranulation profile changed from a bell-shape response curve in WT cells to a threshold response (similar to that of DNP-BSA) in Lyn^KO^ cells. These results show that the structure of antigens can create signaling environments that modulate the recruitment of positive and negative interaction partners. We have also shown previously that altering Syk dynamics can have dramatic effects on signaling outcomes (10,44). In that work, we showed that the presence of SHIP1 in the signalosome reduces the Syk-FcεRI interaction time, leading to reduced Syk and LAT phosphorylation (10). A similar mechanism may occur in the SHIP1-high DF3 aggregates and remains to be explored. Therefore, the combination of receptor phosphorylation states and the interplay between recruited signaling partners work together to tune outcomes.

Geometric modeling of antigen-induced receptor crosslinking predicted that DNP-BSA and DF3 would form aggregates of distinct geometry, with DF3 forming smaller, less complex aggregates than DNP-BSA. The simulations also measured differences in IgE-to-IgE spacing around each antigen, with separations 8.5-15.2 nm for DF3 and 21.7 nm for the furthest hapten spacing on DNP-BSA, as shown in Figure 1. One limitation in the modeling is that the exact location of the haptens on BSA is not known because the labeling of the BSA amine groups is expected to be a Poisson distribution and can vary between BSA molecules. Therefore, while the randomly selected locations on BSA for the 14 DNPs does not incorporate all possible configurations, the modeling provides insight into the potential aggregate geometry.

Previous studies have relied on the known valency and spacing of synthetic antigens to probe the connection between crosslinked receptor spacing and signaling. Here, we used dSTORM super-resolution imaging combined with BaGoL analysis to directly visualize the nanoscale organization of FcεRI aggregates. Consistent with the Monte Carlo simulations, we found that DF3 aggregates are smaller than DNP-BSA aggregates. Measurements of circularity also show that DF3 aggregates are shifted towards a more linear shape. This change in shape can be interpreted as a loss in complexity due to reduced branching as predicted by simulation. Our calculation of NND from the dSTORM imaging was also consistent with the modeling, with receptors in DF3 aggregates being more closely packed than for DNP-BSA. Previous work using rigid DNA rulers had found that optimal spacing between antigen epitopes (< 10 nm) acts to promote signaling (21). More recently, the use of DNA origami to present DNP with controlled valency and spacing concluded that hapten proximity (~ 6 nm) was more important than number of haptens present (27). We were, therefore, surprised to find that DF3 induced aggregates with closer receptors packing (mode of 6.91 nm) than DNP-BSA (mode of ~ 8.5 nm) but was not able to promote signaling as efficiently. One explanation is that the receptors are within the minimal distance for signaling for each antigen and that mechanisms in addition to receptor spacing, such as aggregate size or complexity, are at play in regulating FcεRI signaling.

There are multiple other factors that may contribute to differential signalosome formation. One is that changes in the surrounding lipid environment can drive recruitment of PH-domain-containing and palmitoylated/myristoylated proteins, such as SFKs. Multiple studies have shown that, after crosslinking, FcεRI associates with cholesterol-enriched, liquid-ordered membrane domains (57–61), supporting a role for ordered membrane domains in bringing Lyn to the receptor. The difference in receptor density within aggregates may coordinate the lipid environment, as evidenced by the interplay between protein condensates and lipid organization (62). The aggregate environment may also include or exclude signaling partners due to size or the effects of condensate formation (59,62–64). Along the same line, the smaller and less complex DF3 aggregates may enable easier phosphatase access, possibly explaining the reduction in γ-phosphorylation (64,65) in DF3 aggregates. Further work will be important to determine how membrane lipid organization and recruitment of other signaling partners may be influenced by antigen geometries to establish unique signaling platforms.

In summary, we have combined biochemical assays with computational modeling and advanced imaging methods to connect the strength of FcεRI signaling with the nanoscale organization of receptor aggregates. While previous studies have found a direct relationship between closer antigen spacing and increased signaling, we found that the size and complexity of the aggregates formed also play a role in directing signaling. The use of super-resolution imaging via fluorescently labeled IgE allowed for quantifying aggregate geometry without modification of the antigen. This approach opens the door to future studies of aggregates formed by natural allergens where antigen properties are not necessarily known or controllable.

## Author Contributions

D.A.R., W.K.K., B.S.W., D.S.L. conceived the overall project. D.A.R was responsible for most biochemical experiments, with assistance from H.C.K, M.P.P, R.M.G, S.L.L. W.K.K. and R.M.G. were responsible for the dSTORM studies with assistance from D.A.R and M.J.W. on analysis. EB performed SPT experiments. Super-resolution imaging and BAMF analysis methods were developed by K.A.L. and M.J.W, with imaging performed by W.K.K. D.S.L supervised the overall project. J.C.D. performed the Monte Carlo simulations with oversight by B.J. and L.T. D.S.L, D.A.R., W.K.K and R.M.G. wrote the manuscript, on which all authors commented.

## Acknowledgments

This work was supported by the National Institutes of Health R35GM126934 to DSL, R21GM132716 and 1R01GM140284 to KAL; National Science Foundation IIS-1553266 to LT. MPP was supported by UNM U-RISE 1T34GM145428 01. We thank Dr. Cedric Cleyrat and Eunice Choi for generation of the knockout cell lines and initial degranulation experiments. We thank Danielle Burke for assistance with plasmid generation and Dr. Mara Steinkamp for helpful discussion. We gratefully acknowledge use of the University of New Mexico Cancer Center fluorescence microscopy and flow cytometry facilities, as well as NIH-NCI support via P30CA118100 for these cores. This work was conducted with support from the University of New Mexico Office of the Vice President for Research Program for Enhancing Research Capacity, was supported by grants from NVIDIA and utilized an NVIDIA A6000 GPU. Molecular graphics and analyses performed with UCSF ChimeraX, developed by the Resource for Biocomputing, Visualization, and Informatics at the University of California, San Francisco, with support from National Institutes of Health R01-GM129325 and the Office of Cyber Infrastructure and Computational Biology, National Institute of Allergy and Infectious Diseases. We would like to thank the UNM Center for Advanced Research Computing, supported in part by the National Science Foundation, for providing the high-performance computing resources used in this work. Any opinions, findings, conclusions, recommendations, or views contained in this document are those of the authors and should not be interpreted as representing the official policies, either expressed or implied, of the NSF or U.S. Government. The U.S. Government is authorized to reproduce and distribute reprints for Government purposes notwithstanding any copyright notation herein.

## References

1. Li, Y., P. S. C. Leung, M. E. Gershwin, and J. Song. 2022. New Mechanistic Advances in FcεRI-Mast Cell-Mediated Allergic Signaling. Clin Rev Allergy Immunol. 63(3):431–446, doi: 10.1007/s12016-022-08955-9.

2. Garman, S. C., B. A. Wurzburg, S. S. Tarchevskaya, J. P. Kinet, and T. S. Jardetzky. 2000. Structure of the Fc fragment of human IgE bound to its high-affinity receptor Fc epsilonRI alpha. Nature. 406(6793):259–266, doi: 10.1038/35018500.

3. Kimura, T., H. Sakamoto, E. Appella, and R. P. Siraganian. 1997. The negative signaling molecule SH2 domain-containing inositol-polyphosphate 5-phosphatase (SHIP) binds to the tyrosine-phosphorylated beta subunit of the high affinity IgE receptor. J Biol Chem. 272(21):13991–13996, doi: 10.1074/jbc.272.21.13991.

4. Bruhns, P., F. Vely, O. Malbec, W. H. Fridman, E. Vivier, and M. Daeron. 2000. Molecular basis of the recruitment of the SH2 domain-containing inositol 5-phosphatases SHIP1 and SHIP2 by fcgamma RIIB. J Biol Chem. 275(48):37357–37364, doi: 10.1074/jbc.M003518200.

5. Hořejší, V., W. Zhang, and B. Schraven. 2004. Transmembrane adaptor proteins: organizers of immunoreceptor signalling. Nat Rev Immunol. 4(8):603–616, doi: 10.1038/nri1414.

6. Ivashkiv, L. B. 2009. Cross-regulation of signaling by ITAM-associated receptors. Nat Immunol. 10(4):340–347, doi: 10.1038/ni.1706.

7. Mahajan, A., D. Barua, P. Cutler, D. S. Lidke, F. A. Espinoza, C. Pehlke, R. Grattan, Y. Kawakami, C. S. Tung, A. R. Bradbury, W. S. Hlavacek, and B. S. Wilson. 2014. Optimal aggregation of FcεRI with a structurally defined trivalent ligand overrides negative regulation driven by phosphatases. ACS Chem Biol. 9(7):1508–1519, doi: 10.1021/cb500134t.

8. Huber, M., C. D. Helgason, J. E. Damen, L. Liu, R. K. Humphries, and G. Krystal. 1998. The src homology 2-containing inositol phosphatase (SHIP) is the gatekeeper of mast cell degranulation. Proc Natl Acad Sci U S A. 95(19):11330–11335, doi: 10.1073/pnas.95.19.11330.

9. Furumoto, Y., S. Nunomura, T. Terada, J. Rivera, and C. Ra. 2004. The FcepsilonRIbeta immunoreceptor tyrosine-based activation motif exerts inhibitory control on MAPK and IkappaB kinase phosphorylation and mast cell cytokine production. J Biol Chem. 279(47):49177–49187, doi: 10.1074/jbc.M404730200.

10. Kanagy, W. K., C. Cleyrat, M. Fazel, S. R. Lucero, M. P. Bruchez, K. A. Lidke, B. S. Wilson, and D. S. Lidke. 2022. Docking of Syk to FcεRI is enhanced by Lyn but limited in duration by SHIP1. Mol Biol Cell. 33(10):ar89, doi: 10.1091/mbc.E21-12-0603.

11. Wakefield, D. L., D. Holowka, and B. Baird. 2017. The FcεRI Signaling Cascade and Integrin Trafficking Converge at Patterned Ligand Surfaces. Mol Biol Cell. 28(23):3383–3396, doi: 10.1091/mbc.E17-03-0208.

12. Saitoh, S., R. Arudchandran, T. S. Manetz, W. Zhang, C. L. Sommers, P. E. Love, J. Rivera, and L. E. Samelson. 2000. LAT is essential for Fc(epsilon)RI-mediated mast cell activation. Immunity. 12(5):525–535, doi: 10.1016/s1074-7613(00)80204-6.

13. Lee, T. D., M. Swieter, and A. D. Befus. 1986. Mast cell responses to helminth infection. Parasitol Today. 2(7):186–191, doi: 10.1016/0169-4758(86)90190-0.

14. Di Lorenzo, G., P. Mansueto, M. Melluso, G. Candore, A. Colombo, M. E. Pellitteri, A. Drago, M. Potestio, and C. Caruso. 1997. Allergic rhinitis to grass pollen: measurement of inflammatory mediators of mast cell and eosinophils in native nasal fluid lavage and in serum out of and during pollen season. J Allergy Clin Immunol. 100(6 Pt 1):832–837, doi: 10.1016/s0091-6749(97)70281-1.

15. de Weck, A. L. 1984. Pathophysiologic mechanisms of allergic and pseudo-allergic reactions to foods, food additives and drugs. Ann Allergy. 53(6 Pt 2):583–586.

16. Atherton, D. J. 1984. Diagnosis and management of skin disorders caused by food allergy. Ann Allergy. 53(6 Pt 2):623–628.

17. Galli, S. J., P. Starkl, T. Marichal, and M. Tsai. 2017. Mast Cells and IgE can Enhance Survival During Innate and Acquired Host Responses to Venoms. Trans Am Clin Climatol Assoc. 128:193–221.

18. Mahajan, A., L. A. Youssef, C. Cleyrat, R. Grattan, S. R. Lucero, C. P. Mattison, M. F. Erasmus, B. Jacobson, L. Tapia, W. S. Hlavacek, M. Schuyler, and B. S. Wilson. 2017. Allergen Valency, Dose, and FcεRI Occupancy Set Thresholds for Secretory Responses to Pen a 1 and Motivate Design of Hypoallergens. J Immunol. 198(3):1034–1046, doi: 10.4049/jimmunol.1601334.

19. Gieras, A., B. Linhart, K. H. Roux, M. Dutta, M. Khodoun, D. Zafred, C. R. Cabauatan, C. Lupinek, M. Weber, M. Focke-Tejkl, W. Keller, F. D. Finkelman, and R. Valenta. 2016. IgE epitope proximity determines immune complex shape and effector cell activation capacity. J Allergy Clin Immunol. 137(5):1557–1565, doi: 10.1016/j.jaci.2015.08.055.

20. Holowka, D., D. Sil, C. Torigoe, and B. Baird. 2007. Insights into immunoglobulin E receptor signaling from structurally defined ligands. Immunol Rev. 217:269–279, doi: 10.1111/j.1600-065X.2007.00517.x.

21. Sil, D., J. B. Lee, D. Luo, D. Holowka, and B. Baird. 2007. Trivalent ligands with rigid DNA spacers reveal structural requirements for IgE receptor signaling in RBL mast cells. ACS Chem Biol. 2(10):674–684, doi: 10.1021/cb7001472.

22. Paar, J. M., N. T. Harris, D. Holowka, and B. Baird. 2002. Bivalent ligands with rigid double-stranded DNA spacers reveal structural constraints on signaling by Fc epsilon RI. J Immunol. 169(2):856–864, doi: 10.4049/jimmunol.169.2.856.

23. Andrews, N. L., J. R. Pfeiffer, A. M. Martinez, D. M. Haaland, R. W. Davis, T. Kawakami, J. M. Oliver, B. S. Wilson, and D. S. Lidke. 2009. Small, mobile FcepsilonRI receptor aggregates are signaling competent. Immunity. 31(3):469–479, doi: 10.1016/j.immuni.2009.06.026.

24. Spendier, K., A. Carroll-Portillo, K. A. Lidke, B. S. Wilson, J. A. Timlin, and J. L. Thomas. 2010. Distribution and dynamics of rat basophilic leukemia immunoglobulin E receptors (FcepsilonRI) on planar ligand-presenting surfaces. Biophys J. 99(2):388–397, doi: 10.1016/j.bpj.2010.04.029.

25. Carroll-Portillo, A., K. Spendier, J. Pfeiffer, G. Griffiths, H. Li, K. A. Lidke, J. M. Oliver, D. S. Lidke, J. L. Thomas, B. S. Wilson, and J. A. Timlin. 2010. Formation of a mast cell synapse: Fc epsilon RI membrane dynamics upon binding mobile or immobilized ligands on surfaces. J Immunol. 184(3):1328–1338, doi: 10.4049/jimmunol.0903071.

26. Suzuki, R., S. Leach, W. Liu, E. Ralston, J. Scheffel, W. Zhang, C. A. Lowell, and J. Rivera. 2014. Molecular editing of cellular responses by the high-affinity receptor for IgE. Science. 343(6174):1021–1025, doi: 10.1126/science.1246976.

27. Schneider, L., K. S. Rabe, C. M. Domínguez, and C. M. Niemeyer. 2023. Hapten-Decorated DNA Nanostructures Decipher the Antigen-Mediated Spatial Organization of Antibodies Involved in Mast Cell Activation. ACS Nano. 17(7):6719–6730, doi: 10.1021/acsnano.2c12647.

28. Posner, R. G., D. Geng, S. Haymore, J. Bogert, I. Pecht, A. Licht, and P. B. Savage. 2007. Trivalent antigens for degranulation of mast cells. Org Lett. 9(18):3551–3554, doi: 10.1021/ol071175h.

29. Liu, F. T., J. W. Bohn, E. L. Ferry, H. Yamamoto, C. A. Molinaro, L. A. Sherman, N. R. Klinman, and D. H. Katz. 1980. Monoclonal dinitrophenyl-specific murine IgE antibody: preparation, isolation, and characterization. J Immunol. 124(6):2728–2737.

30. Fazel, M., M. J. Wester, D. J. Schodt, S. R. Cruz, S. Strauss, F. Schueder, T. Schlichthaerle, J. M. Gillette, D. S. Lidke, B. Rieger, R. Jungmann, and K. A. Lidke. 2022. High-precision estimation of emitter positions using Bayesian grouping of localizations. Nat Commun. 13(1):7152, doi: 10.1038/s41467-022-34894-2.

31. Valley, C. C., D. J. Arndt-Jovin, N. Karedla, M. P. Steinkamp, A. I. Chizhik, W. S. Hlavacek, B. S. Wilson, K. A. Lidke, and D. S. Lidke. 2015. Enhanced dimerization drives ligand-independent activity of mutant epidermal growth factor receptor in lung cancer. Mol Biol Cell. 26(22):4087–4099, doi: 10.1091/mbc.E15-05-0269.

32. Schwartz, S. L., Q. Yan, C. A. Telmer, K. A. Lidke, M. P. Bruchez, and D. S. Lidke. 2015. Fluorogen-activating proteins provide tunable labeling densities for tracking FcεRI independent of IgE. ACS Chem Biol. 10(2):539–546, doi: 10.1021/cb5005146.

33. Travers, T., W. K. Kanagy, R. A. Mansbach, E. Jhamba, C. Cleyrat, B. Goldstein, D. S. Lidke, B. S. Wilson, and S. Gnanakaran. 2019. Combinatorial diversity of Syk recruitment driven by its multivalent engagement with FcεRIγ. Mol Biol Cell. 30(17):2331–2347, doi: 10.1091/mbc.E18-11-0722.

34. Metzger, H., G. Alcaraz, R. Hohman, J. P. Kinet, V. Pribluda, and R. Quarto. 1986. The receptor with high affinity for immunoglobulin E. Annu Rev Immunol. 4:419–470, doi: 10.1146/annurev.iy.04.040186.002223.

35. Wilson, B. S., J. R. Pfeiffer, and J. M. Oliver. 2000. Observing FcepsilonRI signaling from the inside of the mast cell membrane. J Cell Biol. 149(5):1131–1142, doi: 10.1083/jcb.149.5.1131.

36. Manavi, K., B. S. Wilson, and L. Tapia (2012). Simulation and analysis of antibody aggregation on cell surfaces using motion planning and graph analysis. Proceedings of the ACM Conference on Bioinformatics, Computational Biology and Biomedicine. Association for Computing Machinery.

37. James, L. C., P. Roversi, and D. S. Tawfik. 2003. Antibody multispecificity mediated by conformational diversity. Science. 299(5611):1362–1367, doi: 10.1126/science.1079731.

38. Leonard, P., P. D. Scotney, T. Jabeen, S. Iyer, L. J. Fabri, A. D. Nash, and K. R. Acharya. 2008. Crystal structure of vascular endothelial growth factor-B in complex with a neutralising antibody Fab fragment. J Mol Biol. 384(5):1203–1217, doi: 10.1016/j.jmb.2008.09.076.

39. Wan, T., R. L. Beavil, S. M. Fabiane, A. J. Beavil, M. K. Sohi, M. Keown, R. J. Young, J. Henry, R. J. Owens, H. J. Gould, and B. J. Sutton. 2002. The crystal structure of IgE Fc reveals an asymmetrically bent conformation. Nat Immunol. 3(7):681–686, doi: 10.1038/ni811.

40. Güthe, S., L. Kapinos, A. Möglich, S. Meier, S. Grzesiek, and T. Kiefhaber. 2004. Very fast folding and association of a trimerization domain from bacteriophage T4 fibritin. J Mol Biol. 337(4):905–915, doi: 10.1016/j.jmb.2004.02.020.

41. Majorek, K. A., P. J. Porebski, A. Dayal, M. D. Zimmerman, K. Jablonska, A. J. Stewart, M. Chruszcz, and W. Minor. 2012. Structural and immunologic characterization of bovine, horse, and rabbit serum albumins. Mol Immunol. 52(3–4):174–182, doi: 10.1016/j.molimm.2012.05.011.

42. Meng, E. C., T. D. Goddard, E. F. Pettersen, G. S. Couch, Z. J. Pearson, J. H. Morris, and T. E. Ferrin. 2023. UCSF ChimeraX: Tools for structure building and analysis. Protein Sci. 32(11):e4792, doi: 10.1002/pro.4792.

43. Pettersen, E. F., T. D. Goddard, C. C. Huang, E. C. Meng, G. S. Couch, T. I. Croll, J. H. Morris, and T. E. Ferrin. 2021. UCSF ChimeraX: Structure visualization for researchers, educators, and developers. Protein Sci. 30(1):70–82, doi: 10.1002/pro.3943.

44. Schwartz, S. L., C. Cleyrat, M. J. Olah, P. K. Relich, G. K. Phillips, W. S. Hlavacek, K. Lidke, B. S. Wilson, and D. S. Lidke. 2017. Differential mast cell outcomes are sensitive to FcεRI-Syk binding kinetics. Mol Biol Cell. 28(23):3397–3414, doi: 10.1091/mbc.E17-06-0350.

45. Fazel, M., M. J. Wester, H. Mazloom-Farsibaf, M. B. M. Meddens, A. S. Eklund, T. Schlichthaerle, F. Schueder, R. Jungmann, and K. A. Lidke. 2019. Bayesian Multiple Emitter Fitting using Reversible Jump Markov Chain Monte Carlo. Sci Rep. 9(1):13791, doi: 10.1038/s41598-019-50232-x.

46. Andrews, N. L., K. A. Lidke, J. R. Pfeiffer, A. R. Burns, B. S. Wilson, J. M. Oliver, and D. S. Lidke. 2008. Actin restricts FcepsilonRI diffusion and facilitates antigen-induced receptor immobilization. Nat Cell Biol. 10(8):955–963, doi: 10.1038/ncb1755.

47. Schodt, D. J., F. Farzam, S. Liu, and K. A. Lidke. 2023. Automated multi-target super-resolution microscopy with trust regions. Biomed Opt Express. 14(1):429–440, doi: 10.1364/boe.477501.

48. Schodt, D. J., M. J. Wester, M. Fazel, S. Khan, H. Mazloom-Farsibaf, S. Pallikkuth, M. B. M. Meddens, F. Farzam, E. A. Burns, W. K. Kanagy, D. A. Rinaldi, E. Jhamba, S. Liu, P. K. Relich, M. J. Olah, S. L. Steinberg, and K. A. Lidke. 2023. SMITE: Single Molecule Imaging Toolbox Extraordinaire (MATLAB). Journal of Open Source Software. 8(90):5563, doi: 10.21105/joss.05563.

49. Smith, C. S., N. Joseph, B. Rieger, and K. A. Lidke. 2010. Fast, single-molecule localization that achieves theoretically minimum uncertainty. Nat Methods. 7(5):373–375, doi: 10.1038/nmeth.1449.

50. Huang, X. 2011. Quasimaximum likelihood estimation of discretely observed diffusions. The Econometrics Journal. 14(2):241–256, doi:10.1111/j.1368-423X.2010.00324.x, https://doi.org/10.1111/j.1368-423X.2010.00324.x.

51. Schodt, D. J., and K. A. Lidke. 2021. Spatiotemporal Clustering of Repeated Super-Resolution Localizations via Linear Assignment Problem. Front Bioinform. 1:724325, doi: 10.3389/fbinf.2021.724325.

52. Wester, M. J., D. J. Schodt, H. Mazloom-Farsibaf, M. Fazel, S. Pallikkuth, and K. A. Lidke. 2021. Robust, fiducial-free drift correction for super-resolution imaging. Sci Rep. 11(1):23672, doi: 10.1038/s41598-021-02850-7.

53. Sander, J., M. Ester, H.-P. Kriegel, and X. Xu. 1998. Density-Based Clustering in Spatial Databases: The Algorithm GDBSCAN and Its Applications. Data Mining and Knowledge Discovery. 2:169–194, doi: 10.1023/A:1009745219419, https://link.springer.com/article/10.1023/A:1009745219419.

54. Daszykowski, M., B. Walczak, and D. L. Massart. 2002. On the optimal partitioning of data with K-means, growing K-means, neural gas, and growing neural gas. J Chem Inf Comput Sci. 42(6):1378–1389, doi: 10.1021/ci020270w.

55. Xu, K., B. Goldstein, D. Holowka, and B. Baird. 1998. Kinetics of multivalent antigen DNP-BSA binding to IgE-Fc epsilon RI in relationship to the stimulated tyrosine phosphorylation of Fc epsilon RI. J Immunol. 160(7):3225–3235.

56. Wilson, B. S., S. L. Steinberg, K. Liederman, J. R. Pfeiffer, Z. Surviladze, J. Zhang, L. E. Samelson, L. H. Yang, P. G. Kotula, and J. M. Oliver. 2004. Markers for detergent-resistant lipid rafts occupy distinct and dynamic domains in native membranes. Mol Biol Cell. 15(6):2580–2592, doi: 10.1091/mbc.e03-08-0574.

57. Davey, A. M., R. P. Walvick, Y. Liu, A. A. Heikal, and E. D. Sheets. 2007. Membrane order and molecular dynamics associated with IgE receptor cross-linking in mast cells. Biophys J. 92(1):343–355, doi: 10.1529/biophysj.106.088815.

58. Davey, A. M., K. M. Krise, E. D. Sheets, and A. A. Heikal. 2008. Molecular perspective of antigen-mediated mast cell signaling. J Biol Chem. 283(11):7117–7127, doi: 10.1074/jbc.M708879200.

59. Bag, N., A. Wagenknecht-Wiesner, A. Lee, S. M. Shi, D. A. Holowka, and B. A. Baird. 2021. Lipid-based and protein-based interactions synergize transmembrane signaling stimulated by antigen clustering of IgE receptors. Proc Natl Acad Sci U S A. 118(35), doi: 10.1073/pnas.2026583118.

60. Wu, M., D. Holowka, H. G. Craighead, and B. Baird. 2004. Visualization of plasma membrane compartmentalization with patterned lipid bilayers. Proc Natl Acad Sci U S A. 101(38):13798–13803, doi: 10.1073/pnas.0403835101.

61. Shelby, S. A., S. L. Veatch, D. A. Holowka, and B. A. Baird. 2016. Functional nanoscale coupling of Lyn kinase with IgE-FcεRI is restricted by the actin cytoskeleton in early antigen-stimulated signaling. Mol Biol Cell. 27(22):3645–3658, doi: 10.1091/mbc.E16-06-0425.

62. Wang, H. Y., S. H. Chan, S. Dey, I. Castello-Serrano, M. K. Rosen, J. A. Ditlev, K. R. Levental, and I. Levental. 2023. Coupling of protein condensates to ordered lipid domains determines functional membrane organization. Sci Adv. 9(17):eadf6205, doi: 10.1126/sciadv.adf6205.

63. Jaqaman, K., and J. A. Ditlev. 2021. Biomolecular condensates in membrane receptor signaling. Curr Opin Cell Biol. 69:48–54, doi: 10.1016/j.ceb.2020.12.006.

64. Felce, J. H., E. Sezgin, M. Wane, H. Brouwer, M. L. Dustin, C. Eggeling, and S. J. Davis. 2018. CD45 exclusion- and cross-linking-based receptor signaling together broaden FcεRI reactivity. Sci Signal. 11(561), doi: 10.1126/scisignal.aat0756.

65. Young, R. M., X. Zheng, D. Holowka, and B. Baird. 2005. Reconstitution of regulated phosphorylation of FcepsilonRI by a lipid raft-excluded protein-tyrosine phosphatase. J Biol Chem. 280(2):1230–1235, doi: 10.1074/jbc.M408339200.

