## Supplemental Figures for "Antigen properties shape organization of FcεRI aggregates to tune mast cell signaling"

Rinaldi, *et al.*

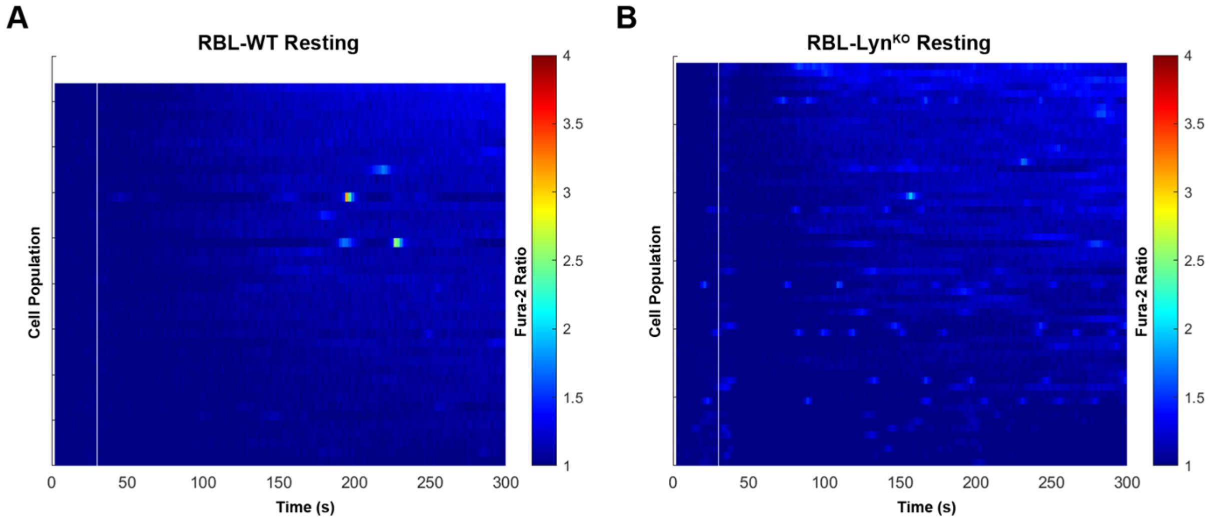


**Figure S1.** Kymographs displaying the Fura-2 ratio over time for resting WT or RBL-Lyn$^{KO}$ (only only HBSS was added at the time point indicated by the white line). Each row represents a single cell trace over time. See also Figure 2.

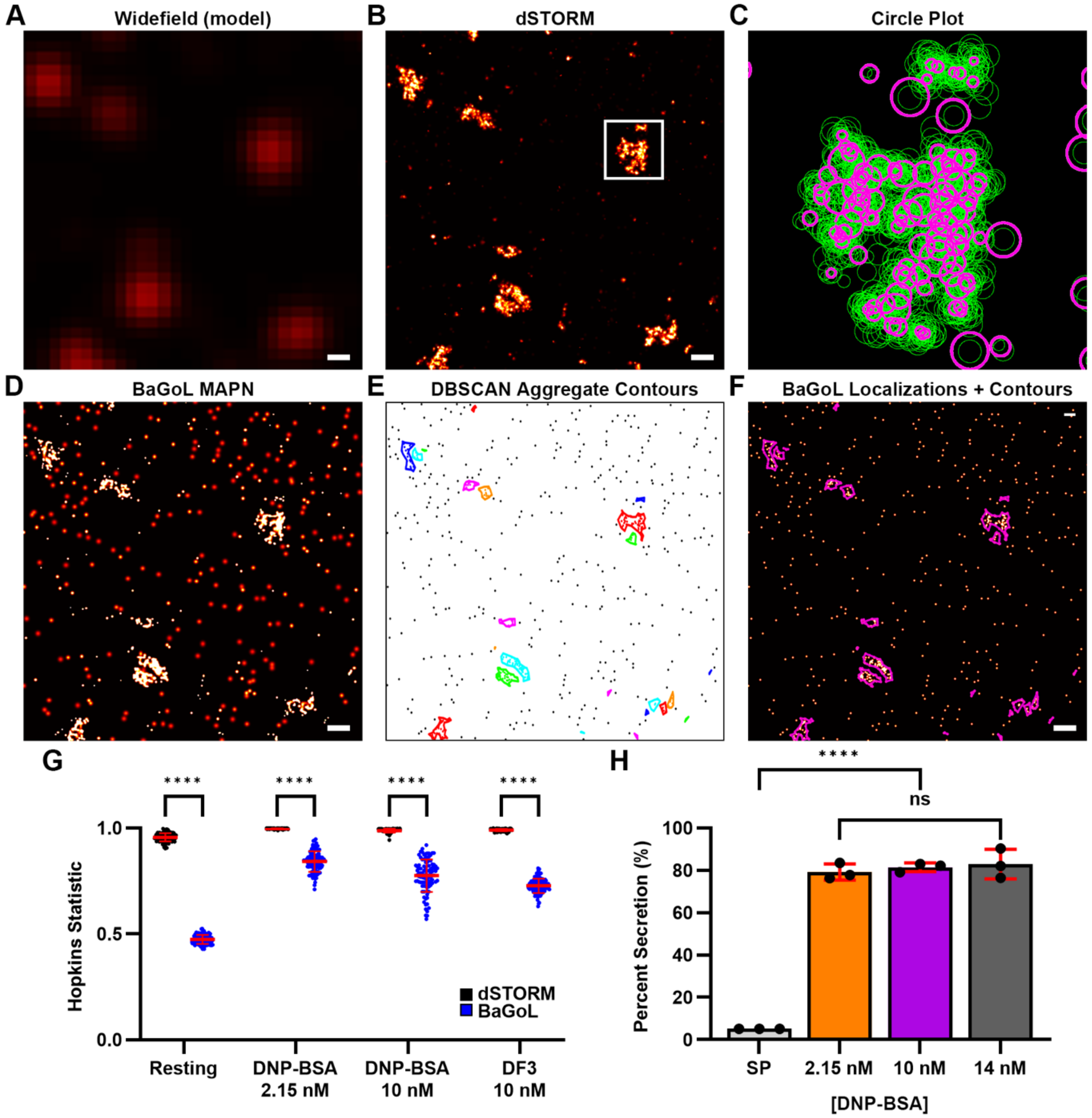


**Figure S2. A-F)** Example of BaGoL analysis workflow. **A)** diffraction limited image; **B)** dSTORM Gaussian image, **C)** zoom in of region (white box) in (**B**) showing BaGoL emitter localizations (magenta) overlayed with dSTORM localizations (green); **D)** post-BaGoL Gaussian image, where Gaussian blobs represent the calculated error for each localization (MAPN); **E)** results of DBSCAN analysis where contours identify aggregates; and **F)** DBSCAN contours over the BaGoL output, where each localization is plotted as 5 nm Gaussian blob to better visualize the individual receptors. Scale bars, 50 nm. **G)** Comparison of Hopkins Statistic between dSTORM and BaGoL images. As expected, Hopkins value is high (near 1) for dSTORM images due to the repeated blinking and reduced in BaGoL collapsed images. Note, BaGoL image values are reproduced here from Fig 7D to directly compare with dSTORM values. **H)** Degranulation response is similar for the three doses of DNP-BSA compared.
